# The Genetic Architecture of Larval Aggregation Behavior in Drosophila

**DOI:** 10.1101/2020.11.01.363994

**Authors:** Ross M McKinney, Yehuda Ben-Shahar

**Affiliations:** Department of Biology, Washington University in St. Louis, St. Louis, MO, USA

**Author notes:** **Contact:** Yehuda Ben-Shahar, Department of Biology, Washington University in St. Louis, St. Louis, MO 63130.

**Keywords:** Drosophila melanogaster, fruit fly, vinegar fly, foraging, sociality

## Abstract

Many insect species exhibit basal social behaviors such as aggregation, which play important roles in their feeding and mating ecologies. However, the evolutionary, genetic, and physiological mechanisms that regulate insect aggregation remain unknown for most species. Here, we used natural populations of Drosophila melanogaster to identify the genetic architecture that drives larval aggregation feeding behavior. By using quantitative and reverse genetic approaches, we have identified a complex neurogenetic network that plays a role in regulating the decision of larvae to feed in either solitude or as a group. Results from single gene, RNAi-knockdown experiments show that several of the identified genes represent key nodes in the genetic network that determines the level of aggregation while feeding. Furthermore, we show that a single non-coding SNP in the gene CG14205, a putative acyltransferase, is associated with both decreased mRNA expression and increased aggregate formation, which suggests that it has a specific role in inhibiting aggregation behavior. Our results identify, for the first time, the genetic components which interact to regulate naturally occurring levels of aggregation in D. melanogaster larvae.

## Introduction

Group formation is one of the simplest forms of social interaction exhibited by individual animals. Yet, the genetic and physiological mechanisms underlying group formation are largely unknown for most species. *Drosophila melanogaster* larvae form simple cooperative group aggregates while feeding, which has been hypothesized to increase their fitness by providing defense against predation, as well as enabling individuals to communally digest food substrates more easily (Prokopy & Roitberg, 2001; Sokolowski, 2010; Wu et al., 2003). Previous studies have suggested that in *Drosophila* and several other insect species, the formation and maintenance of larval aggregation is primarily regulated by the chemosensory detection of aggregation pheromones, as well as other sensory modalities (Leonhardt et al., 2016; Louis & de Polavieja, 2017; Rooke et al., 2020; Steiger & Stokl, 2017; Symonds & Wertheim, 2005; Thibert et al., 2016). Specifically, in *Drosophila melanogaster*, at least two pheromones produced by larvae have been shown to act as chemoattractants (Mast et al., 2014). However, the downstream neural and genetic pathways that regulate larval aggregation behavior remain largely unexplored.

To optimize fitness, the decision of individual larvae on whether to aggregate while feeding is likely regulated by the interplay between attractive and repulsive signals directly emitted by other conspecifics, or indirectly via feeding-related chemical changes of the consumed food. Indeed, it has been shown that food patch choice is influenced by the presence of other larvae, and the decision to choose one food patch over another is a function of group size (Durisko & Dukas, 2013; Lihoreau et al., 2016) and genetics (Allen et al., 2017; Fitzpatrick et al., 2007; Kaun, Hendel, et al., 2007; Kaun, Riedl, et al., 2007). However, although some conserved peptidergic signaling pathways have been shown to regulate aggregation in *Drosophila* larvae (Wu et al., 2003), most signals and downstream neuronal and genetic pathways that regulate group size via attractive and repulsive signals, remain unknown.

Understanding the genetic architecture that underlies insect aggregation is important not only for deciphering the biological principles that drive social decision making in general, but would provide insight into means of offsetting the economic impact of insect pests. To address this important question, we used the *Drosophila* Genetic Reference Panel (DGRP) (Mackay et al., 2012) to identify genetic variations associated with the extent of larval feeding aggregate size. By combining a genome-wide, quantitative genetics approach with single gene manipulations, we have identified several key genes that contribute to group size in natural populations of *Drosophila* larva. Our results highlight the utility of *D. melanogaster* for understanding the genetics of group formation and provide several genetic targets for further research on this topic.

## Materials and Methods

### Animals

All fly lines were reared on standard corn syrup-soy food (Achron Scientific), and kept under a 12h:12h light:dark schedule at 25 °C and 60% humidity. Lines from the Drosophila Genetic Reference Panel (DGRP) (Mackay et al., 2012) used in this study are available from the Bloomington Drosophila Stock Center (BDSC, Bloomington, IN). UAS-RNAi lines and the *elav*- and *tubulin-*GAL4 lines were from either the Bloomington Drosophila Stock Center or the Vienna Drosophila Resource Center (VDRC) (Dietzl et al., 2007; Perkins et al., 2015). All fly lines used in this study, along with their stock numbers and genotypes, are listed in Table S1.

### Larval aggregation assays

Larval aggregation was assayed as follows. Approximately 30, second/third instar larvae were collected from standard vials using a 15% sucrose solution (w/v). Larvae were placed onto the center of a 60mm petri dish containing 20% apple juice (v/v) and 1% agar (w/v) *en masse* and allowed to roam the plate freely for 15 minutes. Subsequently, a picture of the plate was taken (Figure 1A), and the fraction of aggregating larvae was calculated as described below. All behavioral assays were conducted at 25 ^*o*^C and 70 % humidity.

**Figure 1:**
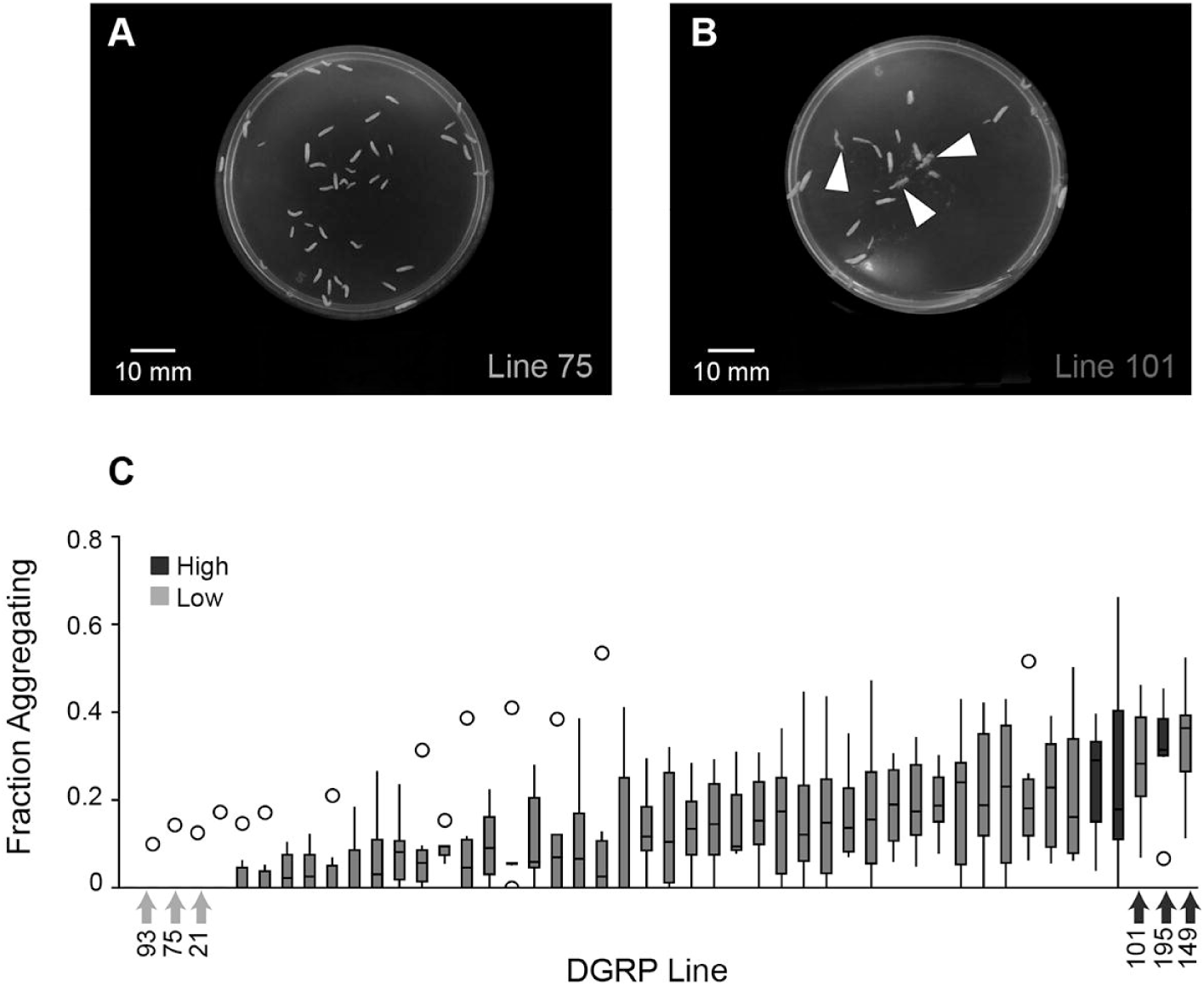
Variation in levels of aggregation between natural populations of Drosophila. **(A)** An image of a DGRP line (Line 75) that showed low levels of aggregation, and **(B)** an image of a DGRP line (Line 101) that showed high levels of aggregation. White arrowheads point to groups of aggregating larvae. **(C)** Boxplots showing the fraction of aggregating larvae for each of the 48 DGRP lines that were included in the GWAS (n=5–9 replicates per line); outliers are shown with open circles. DGRP lines with either low (Low) or high (High) levels of aggregation that were used in subsequent analyses are labeled and shown in either light or dark blue, respectively.

Larval groups were defined as an “aggregate” if two or more larvae were both (i) in physical contact with one another and (ii) burrowing into the agar plate. To calculate the fraction of larvae that were aggregating, we summed the number of larvae forming aggregates and divided it by the total number of larvae observable from the picture taken at the end of the test period.

### Genome Wide Association Study

A total of 4-9 behavioral assays were conducted for each DGRP line, and the mean proportion of aggregating larvae was used for comparison in a genome wide association study (GWAS). A linear regression model was run using the easyGWAS server (Grimm et al., 2017), with default parameters, to search for genotype by phenotype associations. A total of 2,370,987 SNPs from each of 48 DGRP lines were included in the GWAS, after filtering out any SNPs that were of the same genotype across all lines. Linkage disequilibrium and minor allele frequencies (MAF) were calculated using PLINK (Purcell et al., 2007).

### Gene Networks

GeneMANIA was used to predict a functional gene interaction network for all genes identified in the initial GWAS containing SNPs with a p-value of less than 10^−4.5^ (Warde-Farley et al., 2010). A gene was said to contain a SNP if the SNP occurred within ±500 base pairs of its coding exons as annotated in the *Drosophila* reference genome (version 5.57, FB2014 03). Subsequently, co-expression, co-localization, shared protein domains, and protein-protein interactions were used to calculate the gene interaction network, and up to 20 genes that were not identified as significant in the GWAS were allowed to be added to the network. Genes added to the network were selected such they maximized the number of connections between genes already present in the network (Warde-Farley et al., 2010).

### Gene Ontology Analysis

Genes containing SNPs with a p-value of less than 10^−4.5^ were screened for functionally-enriched gene ontologies using the bioprofiling.de servers ProfCom framework (Antonov et al., 2008). All genes included in the functional gene interaction network were also screened for functionally enriched gene ontologies using GeneMANIA (Warde-Farley et al., 2010). The gene interaction network included 20 additional genes that did not contain significant SNPs; the GO terms found to be associated with this network are therefore more general to a set of genes commonly found to interact with one another, rather than those specifically identified in the GWAS.

### Real Time qRT-PCR

mRNA was collected from groups of 10 whole larvae (n=3–4 replicates per line) using Trizol (ThermoFisher) and reverse transcribed to cDNA using SuperScript III Reverse Transcriptase (ThermoFisher). Sybr Green (ThermoFisher) was used to amplify and quantify expression levels for all genes containing significant SNPs identified in the GWAS. Expression values were calculated relative to the *rp49* control gene using the delta delta Ct method, as we have previously described (Hill et al., 2017; Lu et al., 2012; Vernier et al., 2019). All qPCR primers used in this study are listed in Table S2.

### CG14205-GAL4 Transgenic Flies

An approximately 3 kbp (X:19590171–19593107) region of the *CG14205* promoter was synthesized by Integrated DNA Technologies, Inc (IDT) and placed into the pUCIDT-ampR plasmid (IDT). We subcloned this region into the pENTR-1A plasmid (ThermoFisher) using KpnI and XhoI restriction sites on either side of the promoter, and then used Gateway cloning (ThermoFisher) to move the promoter into the pBPGAL4.2::p65 plasmid (Addgene #26229) (Pfeiffer et al., 2010). This plasmid was subsequently injected into BDSC line #24483 (RainbowGene Inc.), and positive offspring were identified and back-crossed into *w*^1118^. The *CG14205*-GAL4 line was crossed with *UAS-mCD8::GFP* (BDSC #32188) and imaged in third-instar larvae.

## Results

### Genetic variation underlying group formation

As *D. melanogaster* larvae develop, they exhibit a gradual increase in aggregation behavior (Wu et al., 2003). However, the overall genetic architecture that drives the quantitative aspects of larval aggregation remains largely unknown. Therefore, to better understand the genetics underlying aggregation, we screened 48 randomly chosen isogenic wild type lines from the *Drosophila* Genetic Reference Panel (DGRP) (Mackay et al., 2012) for levels of aggregation in third instar larvae and subsequently performed a genome wide association study (GWAS) to look for genetic variation associated with this phenotype.

We found that different lines varied significantly in the extent of aggregation, with some lines tending not to form any aggregates (termed “Low” lines) and other lines containing as many as 40-60% of aggregating larvae (termed “High” lines) (Figure 1). We then ran ANOVAs to search for genetic variation (SNPs) associated with the mean fraction of aggregating larvae across lines (Shorter et al., 2015; Swarup et al., 2013). A total of 2,370,987 ANOVAs were run for each unfiltered SNP in the 48 DGRP lines analyzed, which uncovered 58 significant SNPs (*p <* 10^−5^). Subsequently, 17 protein coding genes that fall within 500 bp of these SNPs were further considered as candidate genes that might be playing a role in larval aggregation decisions (Figure 2, Table S3).

**Figure 2:**
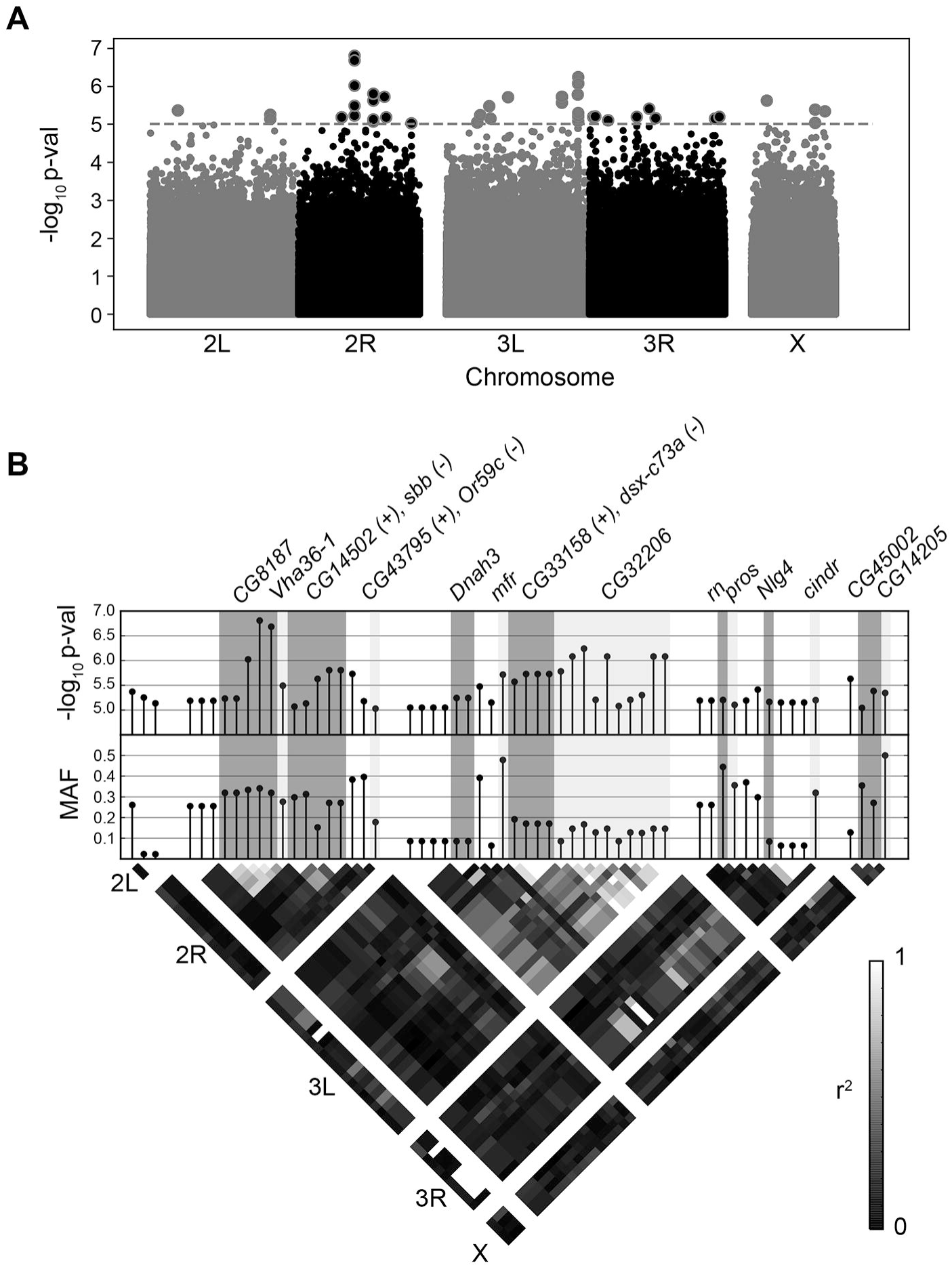
A genome-wide association study identified 58 SNPs that were associated with the extent of larval aggregation across DGRP lines. **(A)** Manhattan plot showing transformed p-values for each of the SNPs included in the GWAS. SNPs with a p-value less than 10−5 (shown by the dashed gray line) were retained for further analysis and are outlined in red. **(B)** A higher resolution view of SNPs highlighted in (A). (Top) Transformed p-values and (Middle) minor allele frequencies (MAFs) for each of the retained SNPs. SNPs that fell within ±500 base pairs of the coding region of a gene are labeled and highlighted together. Some SNPs fell within the coding region of genes on both the plus and minus strand of DNA and are labelled accordingly. (Bottom) Linkage disequilibrium matrix between all of the retained SNPs.

### The neurogenetic network of larval aggregation behavior

To investigate whether specific genetic pathways might be playing a role in larval aggregation decisions, we next used gene ontology (GO) analyses. Because our initial conservative p<10^−5^ significance threshold yielded only 17 protein-coding genes that might be causally associated with levels of aggregation, we used the less conservative threshold of *p<*10^−4.5^, which increased the number of candidate genes to 68. This analysis indicated that this gene list is enriched for the GO terms “Axon guidance” (GO:0007411, p=0.01) and “Plasma membrane” (GO:0005886, p=0.01). To further expand the analysed gene network, we next extended the empirically defined gene network by using the following edges: co-expression, co-localization, shared protein domains, and protein-protein interactions (Supplemental Figure 1A). GO analysis of the extended gene list was still enriched for “Axon guidance”; however, four out of the top six enriched GO terms are neural-tissue specific (Supplemental Figure 1B). Together, these data suggest that at least some of the genetic variations we have identified impact population level phenotypic variations in aggregation decisions via neuronal functions.

### Genetic variations associated with mRNA expression levels

Single nucleotide polymorphisms falling within promoter and enhancer regions of a protein coding gene often affect mRNA expression levels (Khurana et al., 2016; Nord & West, 2020; Visel et al., 2009). Since most of the SNPs we have identified in our GWAS are either intronic or fall upstream of their associated genes (37/46; Table S3), we next tested the hypothesis that some of the identified SNPs affect gene action via their effects on mRNA expression levels. To test this hypothesis, we compared the mRNA expression levels of each of the 17 candidate genes identified in our initial conservative screen between the three phenotypically highest (“High”) and three lowest (“Low”) aggregating DGRP lines (Figure 3A, and B) by using real-time qRT-PCR analyses. We found that at least one SNP (X:19488026) was significantly associated with higher mRNA expression levels of its parent gene, *CG14205*, in all “low” lines relative to all “high” lines (one-way ANOVA; *F*(1,4) = 13.43, *p* = 0.02) (Figure 3). These results suggest that this specific SNP is playing a role in regulating the expression or stability of the *CG14205* mRNAs. The location of this SNP immediately downstream of a predicted splice donor site in the annotated intron 5 of *CG14205* (Figure 3C) suggests that it may affect splicing and/ or stability of the pre-mRNA. Furthermore, we found a significant interaction between *CG14205* expression level and SNP genotype on the levels of aggregation between High and Low lines (two-way ANOVA; *F*(2,3) = 403.3, *p <* 0.01). As *CG14205* expression is significantly higher in Low lines than in High lines, these data suggest that higher expression levels of *CG14205* may reduce aggregation in *D. melanogaster* larvae.

**Figure 3:**
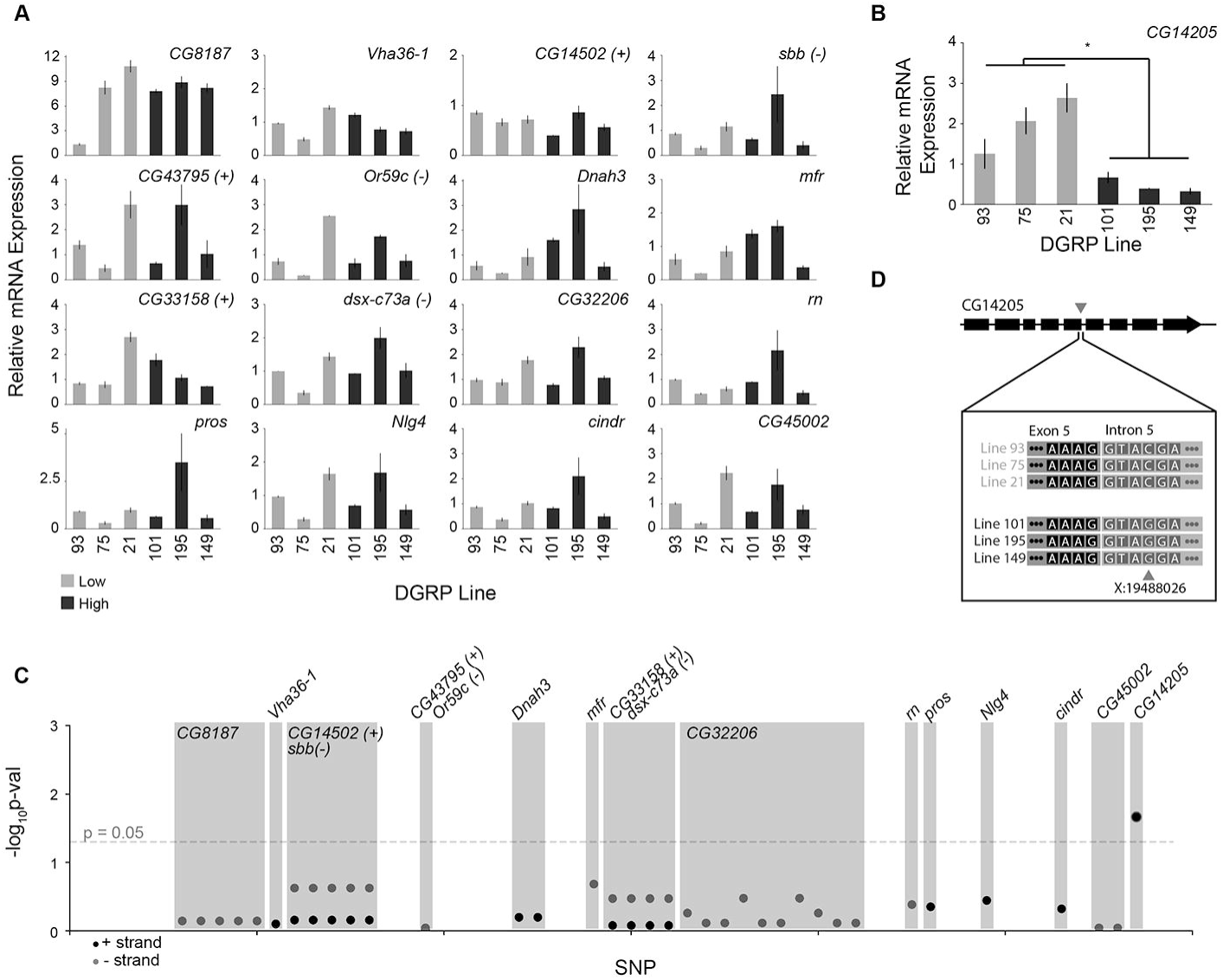
mRNA expression analysis of SNP-containing genes in lines with either low or high levels of aggregation. **(A)** Relative mRNA expression levels for each of the SNP-containing genes identified in the GWAS (n=3–4 replicates per line). Low aggregating lines are shown in light blue, and High aggregating lines are shown in dark blue. **(B)** Relative mRNA expression levels for the *CG14205* gene. A significant association between SNP genotype and *CG14205* mRNA expression was identified (*p <* 0.05; one-way ANOVA), whereby Low aggregating lines had higher levels of expression than High aggregating lines. Note that Low and High lines segregated by genotype, as shown in (C). **(C)** Transformed p-values for associations (ANOVAs) between specific SNP haplotypes and relative mRNA expression level of the gene associated with that SNP. SNPs falling within the same gene are labeled and highlighted together, and SNPs which were significantly associated (*p <* 0.05) with mRNA expression of its gene are outlined in red. **(D)** Genetic architecture of the *CG14205* gene and the DNA sequences surrounding the significantly associated SNP for each of the Low and High DGRP lines. Note that the SNP, X:19488026 (denoted by a red arrow head), falls just past the exon-intron boundary within intron 5 and is positioned to potentially effect mRNA splicing.

Although the biological functions of *CG14205* are unknown, the protein is predicted to be membrane bound Acyltransferase 3 (IPR002656) that is related to the Nose resistant-to-fluoxetine (NRF) protein family in *C. elegans* (Choy & Thomas, 1999). Since several family members have been found to be expressed in the gut epithelium of worms, it has been hypothesized that they may function as novel transporters of lipophilic molecules (Choy et al., 2006). However, the specific biochemical functions of these membrane-bound acyltransferases remain uncharacterized. Nevertheless, previous studies in the moth *Bombyx mori*, have shown that various acyltransferases are required for the synthesis of sex pheromones in moths and other insects (Ding et al., 2016; Mengfang Du et al., 2015; M. Du et al., 2012). Further, a quantitative trait locus (QTL) associated with intra- and interspecific variations in sex pheromones in noctuid moths has been mapped to the regulation of a gene containing a putative Acyltransferase 3 domain (Groot et al., 2013). Therefore, it is possible that *CG14205* plays a direct role in the synthesis of larval aggregation pheromones in *D. melanogaster*.

### Candidate gene knockdown leads to altered levels of aggregation

To further establish a causal role for the genes identified in our initial screen, we studied the effects of neuronal-specific RNAi knockdown of each gene by using the pan-neuronal *elav-GAL4* driver. However, neuronal knockdown of five of the 17 genes we examined (*Vha36-1, dsx-c73a, pros, cindr*, and *CG45002*) was lethal. Of the remaining 12 genes, neuronal knockdown of knockdown of four of the genes (*CG8187, CG14502, CG32206*, and *rn*) lead to higher levels of aggregation relative to controls (Figure 4**A, B**, and **C**). These results suggest that the activity of these four genes affects aggregation decisions in feeding larvae.

**Figure 4:**
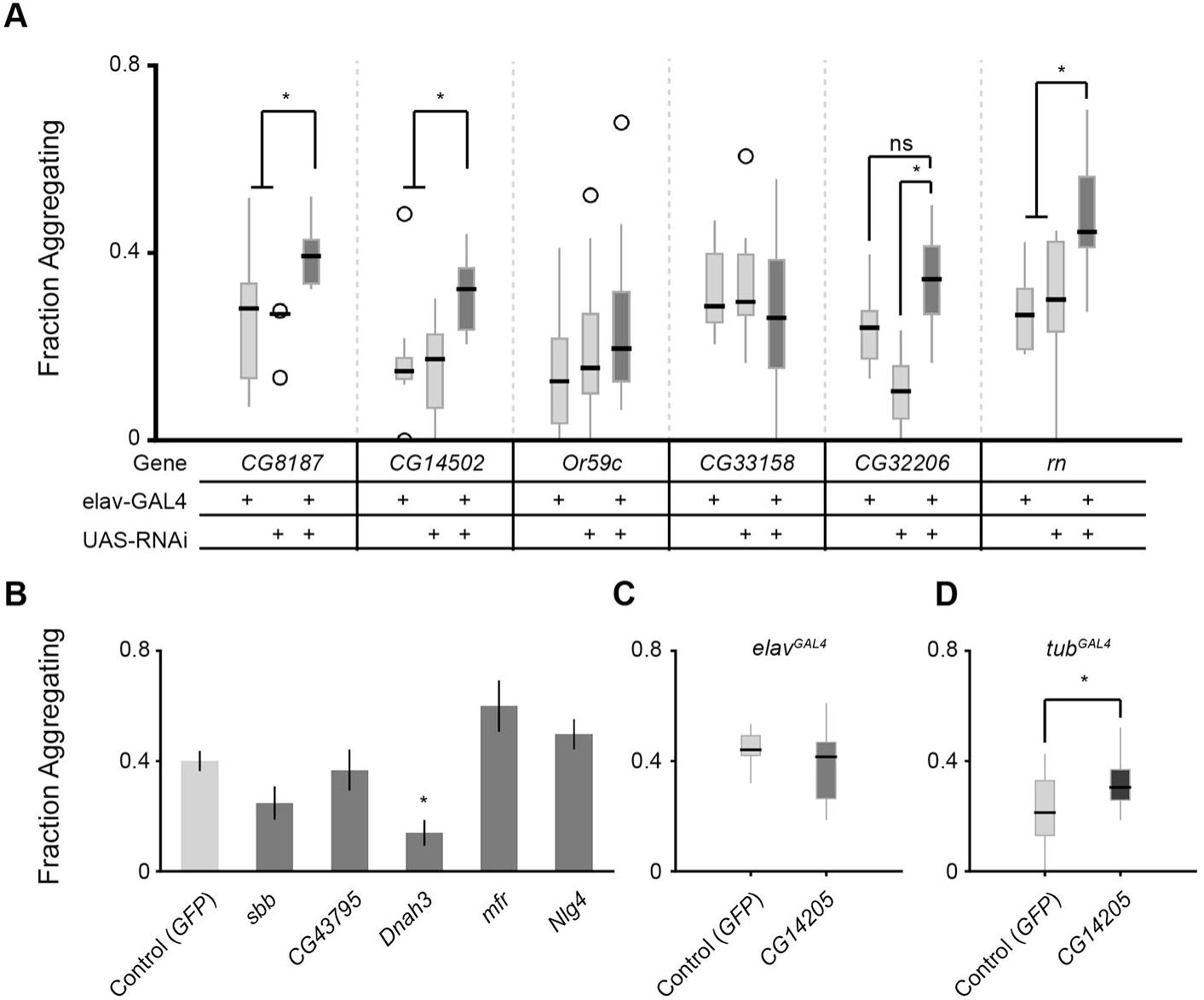
Neuronal knockdown of some candidate genes leads to altered aggregation behavior. **(A)** Pan-neuronal RNAi-mediated knockdown of SNP-associated genes (UAS-RNAi lines from the Vienna Drosophila Resource Center). Knockdown of *CG8187* (n=5-8, *p <* 0.05), *CG14502* (n=8–9, *p <* 0.05), *CG32206* (n=7–9, *p <* 0.01), or *rn* (n=8– 9, *p <* 0.01) using *elav*-GAL4 lead to increased levels of aggregation when compared to parental controls (n=6–19, for all other lines). All statistical comparisons used one-way ANOVA followed by a Tukeys HSD *post-hoc* test. **(B)** Pan-neuronal RNAi-mediated knockdown of SNP-associated genes (UAS-RNAi lines from the Bloomington TRiP collection). Knockdown of *Dnah3* using *elav*-GAL4 lead to a decrease in fraction of larvae aggregating (n=7–17, *p <* 0.01), whereas no other gene knockdowns were significantly different from control (n=4–17). Pairwise Students T-tests were run between each gene knockdown and control to look for statistical significance, and p-values were adjusted for multiple comparisons using a Bonferroni correction. **(C)** TRiP-RNAi-mediated knockdown of *CG14205* in neural tissues, using the *elav*-GAL4 driver, did not lead to altered aggregation (n=8 per group, *p >* 0.05; onetailed Students T-test). **(D)** TRiP-RNAi-mediated knockdown of *CG14205* in all tissues, using the *tubulin*-GAL4 driver, led to a significant increase in the fraction of larvae aggregating compared to control (n=11–12, *p* = 0.025; one-tailed Students T-test).

In contrast, knockdown of *CG14205* in neural tissues did not significantly alter aggregation levels. Given the strong association between the specific *CG14205* alleles, mRNA expression levels, and aggregation levels, we next tested whether genetic variation in this specific gene affect aggregation decision via its action in non-neuronal tissues by using the ubiquitous *tubulin-*GAL4 driver to knockdown *CG14205* in all tissues. As *CG14205* mRNA is expressed to a greater extent in Low aggregating lines, we hypothesized that knocking down *CG14205* should lead to increased levels of aggregation. Indeed, global *CG14205* knockdown resulted in an increase in the fraction of larvae aggregating (one-tailed, Student’s T-test, *p* = 0.025; Figure 4**D**). These results suggest that *CG14205* functions to suppress aggregation in *D. melanogaster* larvae via neuronal-independent signalling pathways in the larval midgut.

While we do not know yet how the midgut activity levels of *CG14205* might affect the decision of individual larvae to join a group, it is likely that this decision controlled by both external sensory stimuli and internal receptors which detect those stimuli. It is possible that the *CG14205* gene is responsible for the biosynthesis or release of a sensory stimulus which inhibits larvae from interacting with one another and forming groups. This hypothesis is consistent with the fact that *CG14205* is required in non-neuronal cells for maintaining normal levels of larval aggregation (compared to controls). Further, mining the FlyExpress and Flygut databases revealed that the expression of *CG14205* is enriched in enterocytes in the larval midgut (Buchon et al., 2013; Celniker et al., 2009) (Figure 5A-B). This expression pattern was further confirmed by imaging the transgenic expression of GFP under the control of the *CG14205* promoter, which revealed strong expression in the most proximal and distal parts of the midgut (Figure 5C-H). Together, these results suggest that *CG14205* plays a role in the synthesis or release, rather than detection, of an inhibitory molecule regulating aggregation.

**Figure 5:**
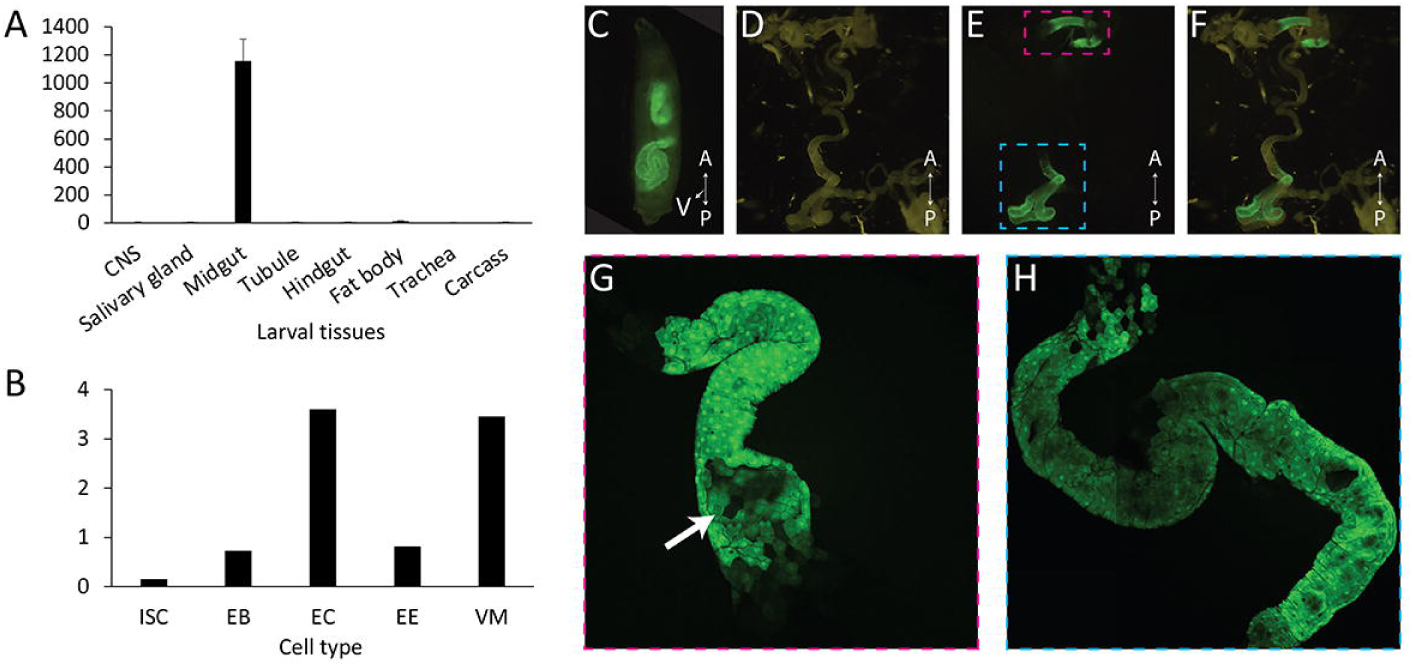
Interaction network of SNP-containing genes. **(A)** Expression levels of *CG14205* across larval tissues. Data were extracted from the FlyAtlas database. **(B)** Expression levels of *CG14205* across midgut cell types. ISC, Intestinal stem cells; EB, Enteroblasts; EC, Enterocytes; EE, Enteroendocrine cells; VM, Visceral muscle. **(C)** *CG14205* expression is restricted to an anterior and posterior regions of the larval midgut. Image of an intact larva expressing GFP under the control of the *CG14205* GAL4 line. **(D-F)** Image of 3^rd^ instar dissected gut: (D) Visible light image, (E) GFP image, (D) Overlay. **(G)** High resolution confocal image of *CG14205*-expressing anterior region (pink dashed box in E). Arrow shows GFP expression in stereotypical enterocytes. **(H)** High resolution confocal image of *CG14205*-expressing posterior region (blue dashed box in E).

## Discussion

It is often assumed that group and social behaviors arise via complex interactions between many genes. Here, we have used an unbiased behavioral quantitative genetic screen to identify population-level natural genetic variations that underlie aggregation in *D. melanogaster* larvae. As expected, our analysis revealed that the decision of individuals on whether to aggregate with other conspecifics is likely depended on a complex genetic network that acts in both neuronal and non-neuronal tissues. Furthermore, by using *in vivo* genetic manipulations, we show that at the population level, both qualitative and quantitative variations could be causally associated with the overall observed behavioral variations between individuals. However, whether the specific identified genes exert their impact on aggregation via a common pathway, and the exact cellular and physiological processes affected by these genes, remain unknown.

Specifically, we found that quantitative expression variations across different alleles of the *CG14205* gene, which encodes a putative acetyl transferase, are strongly associated with larval aggregation while feeding; DGRP lines that exhibit low levels of aggregation express higher levels of *CG14205* transcripts relative to those that display high levels of aggregation (Figure 3B). These data suggest that the activity of *CG14205* inhibits the formation of larval aggregates. While the mechanism regulating this variation in transcript levels is not known, the SNP identified in our initial GWAS screen is adjacent to a predicted intronic splice donor site (Figure 3C), which may affect mRNA splicing and/ or stability via posttranscriptional processes. How *CG14205* activity in the gut might regulate larval aggregation remains unknown. Although our RNAi knockdown studies indicate that *CG14205* is not specifically required in neurons, it remains a possibility that it influences larval behavior via its action in glia or the endocrine system. Alternatively, this gene could be required for the production of a chemical signal that modulates larval aggregation decisions via the enzymatic modification of gut metabolites (Blomquist et al., 2010; Chiu et al., 2019; Hunt & Borden, 1990).

Recent studies have identified both specific chemical cues—pheromones—and receptors to be required for directing aggregation behaviors in *D. melanogaster* larvae (Mast et al., 2014). Although most of what is known about pheromone synthesis in *Drosophila* and other insects relates to cuticular hydrocarbons production by fat-body cells and the oenocytes (Makki et al., 2014; Wicker-Thomas et al., 2015; Zelle et al., 2019), our data indicate that gut derived metabolites can also possibley act as pheromones in *Drosophila*. The possible contribution of *CG14205* to pheromone synthesis is further supported by previous findings about the contribution of acyltransferases to pheromonal signalling in other insect species (Ding et al., 2016; Zhang et al., 2017). Therefore, it is possible that this enzyme functions in the production of some inhibitory chemical cues that *Drosophila* larvae are responsive to during feeding.

Previous studies by us and others have shown that pheromone-driven social interactions in Drosophila and other insects often require the balancing action of both attractive and repulsive cues (Allison & Cardé, 2016; Ben-Shahar et al., 2010; Blomquist & Vogt, 2003; Lu et al., 2012; Lu et al., 2014; McKinney et al., 2015; Zelle et al., 2019). However, in our study, the knockdown of all identified candidate genes leads to increased levels of larval aggregation, which suggest that the primary contributions of these genes are to suppression of aggregation. One possible interpretation of these data is that in natural populations of *D. melanogaster*, it may be that it is more beneficial for larvae to supress aggregation as a function of density to maximize larval fitness. Another non-mutually exclusive explanation might be that our lab assay conditions, and the specific behavioral paradigm used, biased our screen towards the identification of genes whose role contributes specifically to the suppression of larval aggregation.

Nevertheless, our study has uncovered several novel genes involved in directing social aggregation while feeding in *Drosophila* larvae. Although we do not know yet the specific molecular and cellular mechanisms by which any of these genes affect larval feeding behaviors, our data further indicate that natural genetic polymorphisms affect larval social feeding behaviors via both neuronal and non-neuronal pathways (Allen et al., 2017; Anreiter et al., 2017; Sokolowski, 2010).

