## Supplementary material for "The Genetic Architecture of Larval Aggregation Behavior in Drosophila": Supp Fig. 1

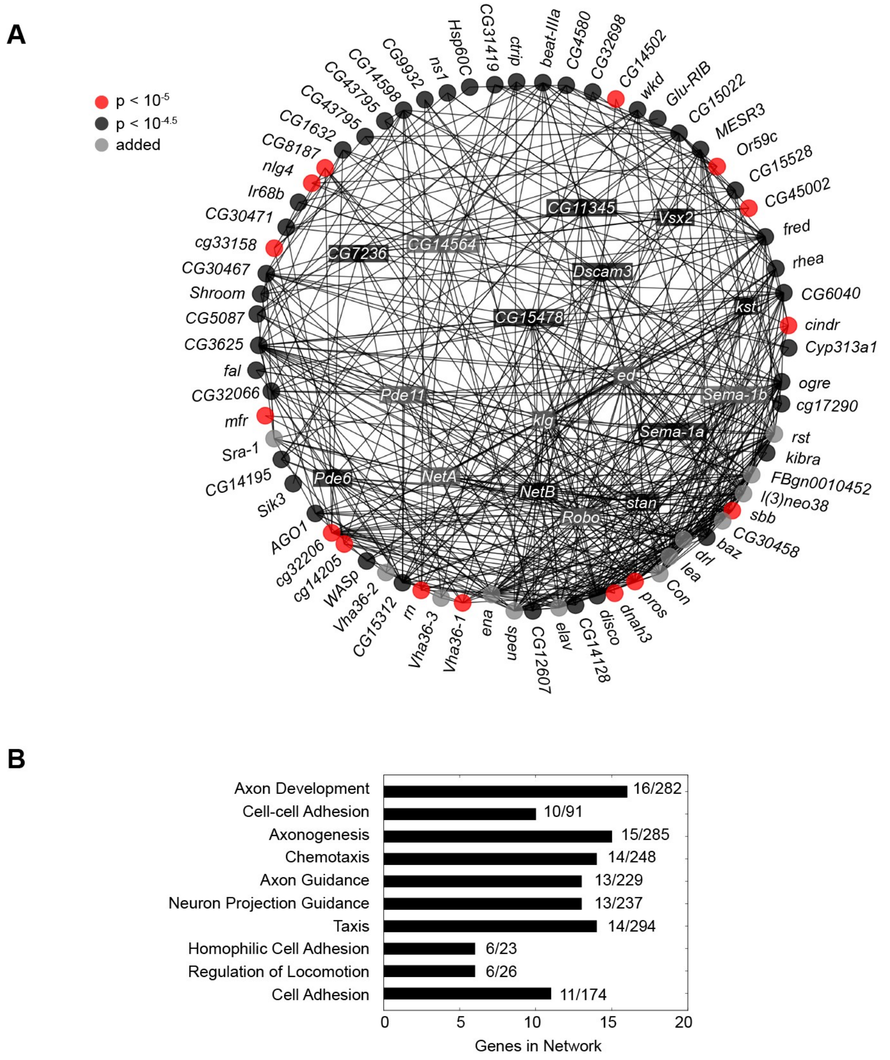


**Supplemental Figure 1:** Interaction network of SNP-containing genes. **(A)** Genes containing SNPs identified in the GWAS at two significance levels ($p < 10^{-4.5}$ [black rectangles/circles] and $p < 10^{-5}$ [red circles]) were used to create a gene interaction network. Red and black circles/rectangles (nodes) in the network represent SNP-containing genes, whereas gray nodes represent genes which were allowed to be added to the network based on the extent of connectivity with SNP-containing genes (see Methods). Lines (edges) connecting each of the nodes represent any of the following interactions: (1) co-expression, (2) co-localization, (3) shared protein domains, (4) and physical protein-protein interactions. **(B)** Enriched functional gene ontologies for all genes present in the network shown in (A).
